# Augmenting Quadriplegic Hand Function Using a Sensorimotor Demultiplexing Neural Interface

**DOI:** 10.1101/604108

**Authors:** PD Ganzer, SC Colachis, MA Schwemmer, DA Friedenberg, CE Swiftney, AF Jacobowitz, DJ Weber, MA Bockbrader, G Sharma

**Affiliations:** Medical Devices and Neuromodulation, Battelle Memorial Institute, 505 King Ave, Columbus, OH, USA 43201; Center for Neuromodulation, The Ohio State University, Columbus, Ohio, 43210, USA.; Department of Physical Medicine and Rehabilitation, The Ohio State University, Columbus, Ohio, 43210, USA.; Advanced Analytics and Health Research, Battelle Memorial Institute, 505 King Ave, Columbus, OH, USA 43201; Department of Bioengineering, University of Pittsburgh, 4200 Fifth Ave, Pittsburgh, PA USA 15260

## Abstract

**Background:** The sense of touch is a key component of motor function. Severe spinal cord injury (SCI) should essentially eliminate sensory information transmission to the brain, that originates from skin innervated from below the lesion. We assessed the hypothesis that, following SCI, residual hand sensory information is transmitted to the brain, can be decoded amongst competing sensorimotor signals, and used to enhance the sense of touch via an intracortically controlled closed-loop brain-computer interface (BCI) system.

**Methods:** Experiments were performed with a participant who has an AIS-A C5 SCI and an intracortical recording array implanted in left primary motor cortex (M1). Sensory stimulation and standard clinical sensorimotor functional assessments were used throughout a series of several mechanistic experiments.

**Findings:** Our results demonstrate that residual afferent hand sensory signals surprisingly reach human primary motor cortex and can be simultaneously demultiplexed from ongoing efferent motor intention, enabling closed-loop sensory feedback during brain-computer interface (BCI) operation. The closed-loop sensory feedback system was able to detect residual sensory signals from up to the C8 spinal level. Using the closed-loop sensory feedback system enabled significantly enhanced object touch detection, sense of agency, movement speed, and other sensorimotor functions.

**Interpretation:** To our knowledge, this is the first demonstration of simultaneously decoding multiplexed afferent and efferent activity from human cortex to control multiple assistive devices, constituting a ‘sensorimotor demultiplexing’ BCI. Overall, our results support the hypothesis that sub-perceptual neural signals can be decoded reliably and transformed to conscious perception, significantly augmenting function.

**Funding:** Internal funding was provided for this study from Battelle Memorial Institute and The Ohio State University Center for Neuromodulation.

## Introduction

Spinal cord injury (SCI) damages sensorimotor circuits leading to paralysis, an impaired sense of agency, and sensory dysfunction. Several studies have employed a brain-computer interface (BCI) to restore motor control via a robotic limb or other assistive device^1–3^. Recent work demonstrates that a once paralyzed limb can be reanimated using motor intention decoded from primary motor cortex (M1)^4–8^, addressing the need of patients with SCI to regain use of their own hand^9^. This decoded M1 signal activates functional electrical stimulation (FES) of the arm musculature to produce the intended hand movement. Unfortunately, ascending sensory information is also disrupted by SCI from key regions of the hand during movement. This further impacts function, as the sense of touch is critical for multiple aspects of motor control^10^. The vast majority of BCI systems do not address these debilitating sensory deficits, ultimately limiting their utility as an interface that addresses both the affected motor and sensory circuits impacted by SCI.

Sensory function can potentially be augmented using a BCI that can decipher residual sensory neural activity from the impaired hand and dynamically translate this into closed-loop sensory feedback that the user can perceive. Intriguingly, recent study demonstrates that residual sub-perceptual sensory information from below the lesion is transmitted to sensory areas of the brain, even following severe clinically complete SCI^11^. M1 may encode sensory and touch-related signals following severe clinically complete SCI, although it has not yet been reported. Essentially all BCI studies that target M1 due so to decode motor intention^1–8^. Sensory information may also be available in M1 that can be used for BCI control.

Although promising, decoding sensory information for closed-loop BCI purposes is challenging for several reasons. First, there is a need to assess how sensory information transmitted from the impaired hand may be represented in the brain, if at all, following severe SCI. Furthermore, sensory and motor neural signals may be multiplexed^12^ together at the BCI recording site, significantly complicating the simultaneous and reliable decoding of multiple device control signals.

## Methods

### Experimental Design

The participant was a 27-year-old male with an AIS-A C5 SCI (with a zone of partial preservation at C6; see Methods, *Study Participant*) and has participated in our previous studies^4–7^. We performed a series of experiments using either passive sensory stimulation (Fig. 1, Supplemental Fig. S1-S4) or active object touch (Fig. 2, 3, Supplemental Fig. S5 & S6) to assess cortical neurophysiology, neural signal decoding, and assistive device control for upper limb functional improvement. Approval for this study was obtained from the U.S. Food and Drug Administration (Investigational Device Exemption) and the Ohio State University Medical Center Institutional Review Board (Columbus, Ohio). The study met institutional requirements for the conduct of human subjects and was registered on the ClinicalTrials.gov website (Identifier NCT01997125; Date: November 22, 2013). All experiments were performed in accordance with the relevant guidelines and regulations set by the Ohio State University Medical Center. The participant referenced in this work provided permission for photographs and videos and completed an informed consent process prior to commencement of the study. Please see the Supplemental Appendix for additional methods details.

**Figure 1.**
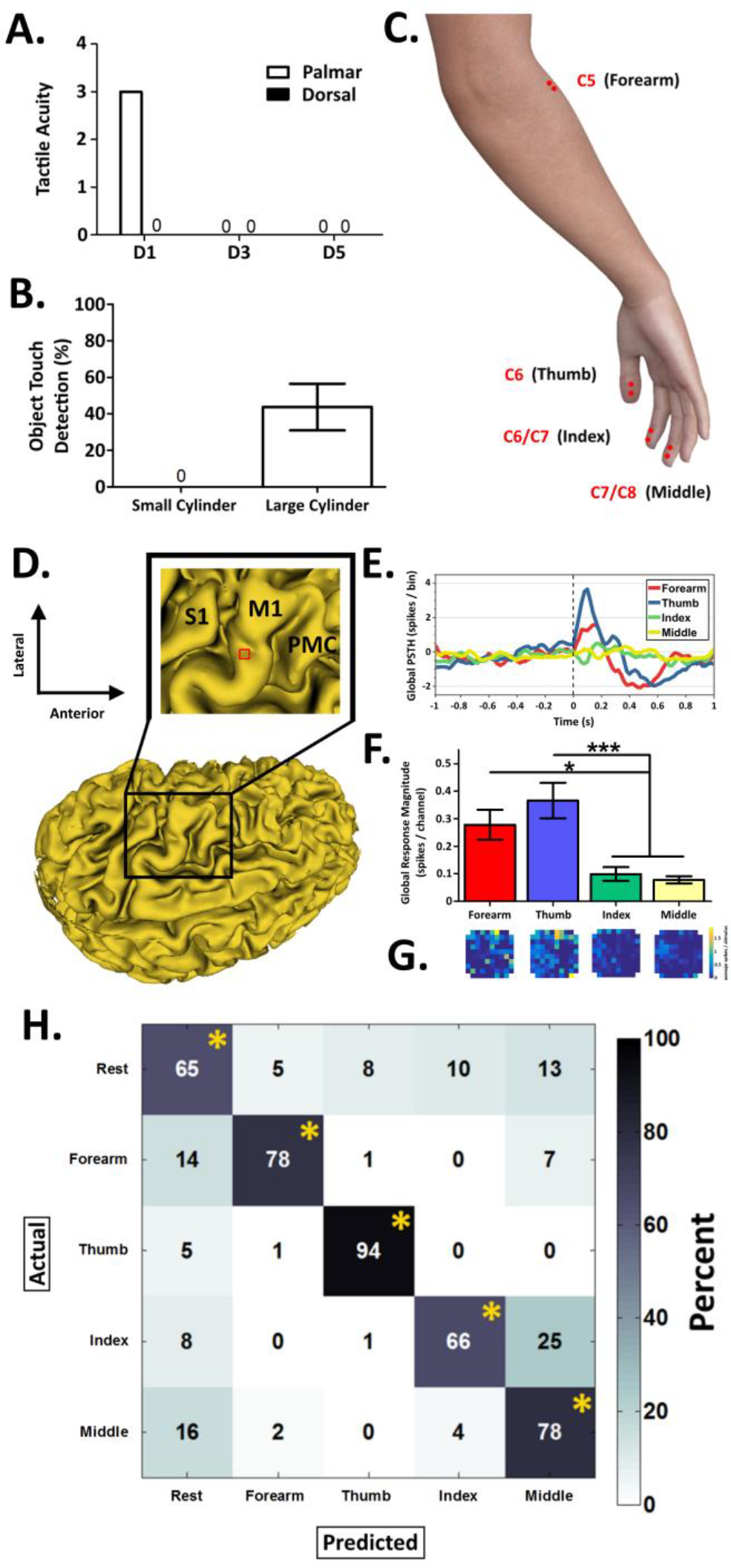
Skin Stimulation on the Arm and Insensate Hand Evokes Robust & Decodable Neural Responses in Contralateral Primary Motor Cortex (M1) Following Cervical Spinal Cord Injury (SCI). **A.** The participant’s hand sensory function was significantly impaired following injury (using standard monofilament testing^32^). **B.** During FES mediated grip, the participant was largely unable to discriminate object touch events in the absence of visual feedback, operating below chance for both standardized objects. These results demonstrate that hand sensory function is dramatically impaired following SCI. **C.** Electrotactile stimulation was performed at 4 different skin locations on the right arm across a period of ~5 months to assess sensory evoked responses in M1. **D.** Pseudocolored 3D reconstruction of the participant’s cerebrum using T1 magnetic resonance imaging. Red box depicts the microelectrode recording array implanted in left M1 (S1 = primary somatosensory cortex; PMC = premotor cortex). **E.** A peri-stimulus time histogram (PSTH) was used to quantify neural modulation in M1 (skin stimulation occurs at time 0, vertical dashed line). Stimulation of the forearm or thumb evoked time locked multiunit activation, with smaller neural responses from index or middle. **F.** Stimulation of the forearm and thumb evoked significantly larger global response magnitudes compared to index or middle (* = p < 0⋅05; *** = p < 0⋅001). **G.** Color coded representations of multiunit response magnitudes across the microelectrode recording array are shown below the labels in panel **C** for each stimulation location (color scaling: blues = no or small neural responses, yellow = large neural responses; units: average spikes / stimulus). **H.** Support vector machine (SVM) decoders were built using neural activity recorded during stimulation at a given skin location or for a rest period (see *Decoding Passive Sensory Stimulation* of the Methods). These passive sensory decoders reliably classified sensory stimulus location, demonstrating significant sensitivity above chance with low false positive rates (confusion matrices show color coded decoder response values, units = percent; * = significantly above chance at p<0⋅001). These results support the hypothesis that somatosensory stimuli evoke decodable neural modulation in contralateral M1 following cervical SCI (data in **D** - **H** is for the maximum stimulation intensity; see Supplemental Fig. S2 for corresponding data at the minimum stimulation intensity). See *Peri-Stimulus Time Histograms* section of the Methods for additional data processing details. Data presented are mean ± S.E.M.

**Figure 2.**
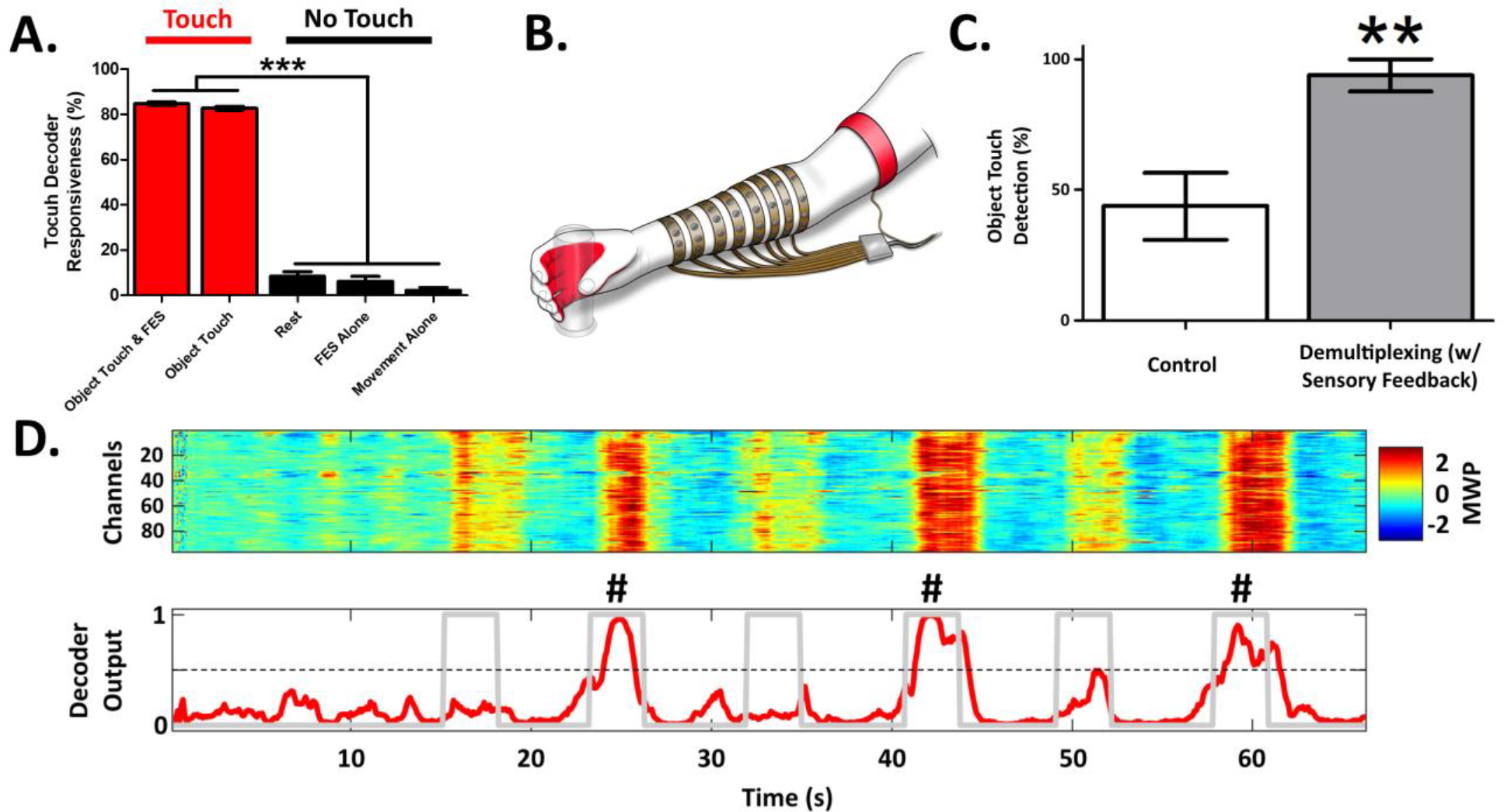
Active Object Touch Can Be Decoded from M1 to Control Closed-Loop Sensory Feedback and Enhance Hand Sensory Function. **A.** Touch decoders were first assessed using ‘Touch’ or ‘No Touch’ periods (see *Decoding Active Object Touch* of the Methods for more details). Touch decoders had significantly higher responsiveness during object touch events (red), compared control cues lacking object touch (black) or rest. Touch decoder false positive rates during cues (data not shown): Object Touch & FES = 12⋅2%; Object Touch = 13⋅7%; FES Alone = 3⋅7%; Movement Alone = 3⋅3%. These results support the hypothesis that machine learning algorithms can be trained to reliably demultiplex active object touch activity from M1. **B.** Touch decoders next controlled closed-loop sensory feedback via a vibrotactile array interfaced with the sensate skin over the ipsilateral bicep (red band in the cartoon schematic). Closed-loop sensory feedback triggered by residual sensory information in M1 more than doubled object touch detection during object grip (**C**, up to ~93%) (** = p < 0⋅01). **D.** Exemplary color-coded mean wavelet power (MWP) input (top) and touch decoder outputs (bottom) during the object touch detection assessment (object placed on cue numbers 2, 4, & 6, # symbol added; cue periods = gray lines; device activation threshold = horizontal dashed line). These results demonstrate that residual sub-perceptual sensory information can be demultiplexed from M1 to trigger closed-loop tactile feedback and significantly improve sensory function. Data presented are mean ± S.E.M.

**Figure 3.**
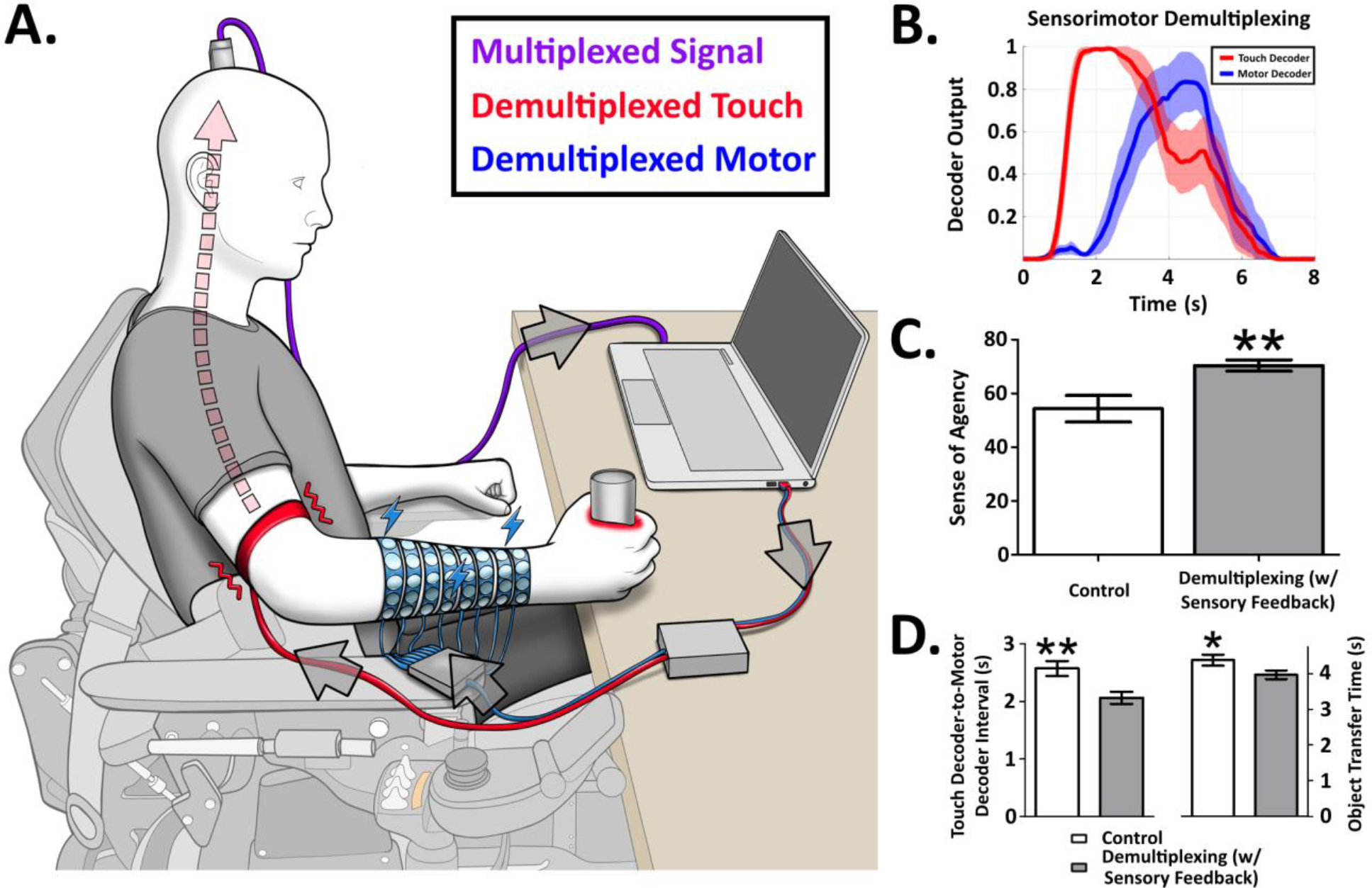
Sensory and Motor Events in M1 Can Be Simultaneously Decoded to Enable ‘Sensorimotor Demultiplexing’ BCI Control and Enhancement of Sensorimotor Function. **A.** Schematic of the participant performing a modified GRT^13^ task with the ‘sensorimotor demultiplexing’ BCI. **B.** We first challenged the touch decoder with a competing simultaneous motor decoder (during a modified GRT). As expected, touch decoders were activated before motor decoders on all object transfers (time 0 = touch cue, followed by participant-initiated motor intention). These results support the hypothesis that afferent touch and efferent motor intention can be simultaneously demultiplexed from M1 during upper limb activity. Closed-loop sensory feedback triggered by demultiplexed sensory neural activity significantly improved the participant’s sense of agency (**C**), motor decoder latency (**D**, left), and object transfer time (**D**, right) (average number of objects transferred per GRT assessment block: control = 9, demultiplexing with sensory feedback = 9⋅75). These results demonstrate the ability to simultaneously decode afferent and efferent information from M1 and activate multiple assistive devices for augmenting sensorimotor function, constituting a ‘sensorimotor demultiplexing’ BCI (* = p < 0⋅05; ** = p < 0⋅01). Data presented are mean ± S.E.M.

## Results

In this study, we sought to perform sensorimotor signal demultiplexing from M1, to enable a BCI system capable of simultaneously controlling multiple devices for enhancing both motor and sensory hand function. All experiments were performed in a chronically paralyzed participant with an AIS-A C5 SCI. We first assessed the participant’s residual hand sensory function (in the absence of visual feedback^26^). He was unable to perceive sensory stimuli to skin innervated below spinal level C6 (clinical tactile assay: Fig. 1A). This sensory impairment was also present during FES-mediated object grip. For example, the participant operated below chance when asked to report if he was gripping an object in the absence of visual feedback (Fig. 1B), a significant sensory impairment further contributing to motor dysfunction.

We next investigated whether residual sensory information could significantly modulate neural activity following skin stimulation (Fig. 1C, Supplemental Fig. S1A-C). Sensory stimuli robustly modulated contralateral M1 (Fig. 1D-G; Supplemental Fig. S2). Stimulation of skin innervated from above or at the C5 SCI evoked time-locked neural modulation, lasting ~10 times longer than the stimulus duration (Fig. 1E). Stimuli applied to skin innervated from below the SCI (index and middle) evoked modest neural modulation in M1, with stimulation to the forearm and thumb evoking significantly larger responses (Fig. 1F: F[3,380] = 9⋅8, p < 0⋅001, Fig. 1G). As expected, separate control experiments demonstrated little to no M1 activation following sensory stimuli to the left arm, ipsilateral to the recording array (Supplemental Fig. S3). These results support the hypothesis that sensory stimuli to skin innervated from both above and below the SCI significantly modulates M1.

Next, we explored whether this sensory activity can be decoded from M1. Decodable sensory events could control a feedback device for improving the impaired sense of touch. We trained a support vector machine (SVM) to detect the skin region being passively stimulated (i.e., a ‘passive sensory decoder’), given the underlying neural activity. Sensory stimulus location was reliably decoded from M1 across a period of several months, performing significantly above chance with low false positive rates (Fig. 1H; Supplemental Fig. S2C). Interestingly, passive sensory decoders for locations that the participant can feel (forearm and thumb) performed equivalently to passive sensory decoders for locations that the participant largely cannot feel (index and middle) (Supplemental Fig. S4). This result demonstrates the ability to decode residual sensory neural activity from M1 that is below conscious perception, from functionally relevant hand dermatomes.

Residual sensory activity could also be decoded in a more challenging context during active object touch using a separate SVM (i.e., a ‘touch decoder’, Fig. 2; see Methods). During validation experiments, touch decoder activation was synchronized to force application from the hand (Supplemental Fig. S5) and performed with high responsiveness during object touch events (~84 %; Fig. 2A, ‘Touch’; F[4,85] = 777, p < 0⋅001; Supplemental Video 1, panel A). As expected, the touch decoder was not activated during control cues which did not have touch events (Fig. 2A, ‘No Touch’; Supplemental Video 1, panels B & C; ‘Rest’ occurs when the cued period is off). These results reveal that residual sensory neural activity can be decoded reliably from M1 during active object manipulation.

The participant was next interfaced with a vibrotactile array on the right bicep, to enable closed-loop sensory feedback (Fig. 2B). This interface was controlled in real time by a touch decoder, to enhance the perception of hand sensory events that are significantly impaired following SCI. The closed-loop sensory feedback system was able to detect residual sensory signals from up to the C8 spinal level, therefore including the insensate regions of the hand (using clinical tactile assay of hand dermatomes; see Methods). Without using the closed-loop feedback system, the participant operated below chance when asked to report if he was gripping an object (Fig. 2C, white; in the absence of visual feedback^26^), largely due to being completely insensate on the vast majority of his hand (clinical tactile assay: Fig. 1A). Closed-loop sensory feedback enabled improved object touch detection from below chance to an over 90% detection rate (Fig. 2C, gray; t(30) = 3⋅5, p = 0⋅001; Fig. 2D) compared to control (Fig. 2C, white). These significant sensory improvements were enabled by sub-perceptual sensory neural activity that is demultiplexed from M1 and enhanced into conscious perception.

Our final set of experiments assessed the hypothesis that afferent and efferent activity in M1 can be demultiplexed to simultaneously control devices for sensory feedback and FES, constituting a ‘sensorimotor demultiplexing’ BCI. The ‘sensorimotor demultiplexing’ BCI system is shown in Fig. 3A. The touch decoder is used to control closed-loop vibrotactile sensory feedback (red band on bicep) and enhance hand touch events. The motor decoder is used to simultaneously control FES of the arm (blue bands on forearm) and produce the desired hand movement. Real-time ‘sensorimotor demultiplexing’ was first demonstrated during a modified grasp and release test (GRT)^13^. The participant cannot perform this task without using the system (data not shown), similar to our previous studies^6,7^. The participant was first cued to position his hand around the object (Fig. 3B, cue at 0 s), and then generate motor intention to activate FES and transfer the object. The touch decoder always preceded the motor decoder, and was time locked to object touch (Fig. 3B; Supplemental Video 2).

We finally enabled ‘sensorimotor demultiplexing’ BCI control using the simultaneous decoding of touch events and motor intention during a set of upper limb assessments. This closed-loop demultiplexing BCI system enabled significant improvements in sense of agency (Fig. 3C; t(46) = 3, p = 0⋅004), motor decoder latency (Fig. 3D, left; t(148) = 2⋅9, p = 0⋅003), and object transfer time (Fig. 3D, right; t(148) = 2⋅1, p = 0⋅03) (Supplemental Fig. S6: exemplary decoder inputs and outputs), compared to a motor-only BCI control. Therefore, rapid closed-loop sensory feedback not only augments sensory function, but also augments motor function. Furthermore, these results provide substantial evidence that sensory feedback during movement can enhance the sense of agency and other benefits of enhanced sensorimotor integration in patients with upper limb dysfunction^14–16^. Overall, successful ‘sensorimotor demultiplexing’ occurred on 100% of task trials (198 total trials). To our knowledge, these findings represent the first demonstration of a BCI system that simultaneously demultiplexes afferent and efferent activity from human cortex for controlling multiple assistive devices and enhancing function.

## Discussion

Severe AIS-A SCI should essentially eliminate sensory information transmission to the brain, that originates from skin innervated from below the lesion. Our results demonstrate that hand sensory signals surprisingly reach M1, even after AIS-A SCI. The participant’s severe C5 SCI functionally blocks communication with the hand, but a clinically complete SCI does not necessarily equate to an anatomically complete SCI. Recent study demonstrates that residual sub-perceptual sensory information from below the lesion is transmitted to sensory areas of the brain, even following severe clinically complete SCI^11^. Our findings support the hypothesis that there is some anatomical sparing of spinal tissue even after severe AIS-A SCI, allowing sensory information transmission from below the lesion to M1, at sufficient levels for enabling new sensory related BCI capabilities. There are likely additional signal types encoded in M1, beyond motor intention and touch related sensory information. In the future, we hope this new set of findings will enable patients with an implanted BCI to maximize the information encoded in the recorded neural activity for new functional gains.

The function restored to the participant using the sensorimotor demultiplexing BCI was significant in several sensorimotor functional domains, ranging from the cognitive control of hand function^18^ to sensorimotor integration. Our control condition throughout all functional assessments was the participant operating the BCI system using only motor control. This control condition is essentially the most challenging control condition possible under the current experimental design. We therefore have designed all experiments to maximally challenge any measured functional improvements during sensorimotor demultiplexing BCI control. Nonetheless, the effect sizes are robust for the functional gains during sensorimotor demultiplexing BCI control. These gains range from an over a 100% improvement to a significant ~0⋅5 second increase in BCI system speed and subsequent upper limb sensorimotor capability. We hypothesize that even larger improvements can be achieved when assessing additional functions in the absence of visual feedback. Closed-loop tactile feedback may mitigate the reliance of BCI users on visually attending to the state of the hand during movement. This would free the user’s visual stream for other important functions during upper limb activity. For example, closed-loop tactile feedback should enable increased BCI operation safety during multitasking, via notifying the user that an object has slipped from their grasp or enabling visual attention to other stimuli, other than the hand, during activities of daily living.

These set of experiments demonstrate the ability to simultaneously reanimate both motor and sensory function in a paralyzed and largely insensate limb. There are alternative ways to provide sensory feedback, including intracortical microsimulation (ICMS) in S1^16,17^. Compared to ICMS in S1, tactile-based feedback enables rapid sensory perception at a significantly faster latency^17^. This was a significant contributor to the choice of using vibrotactile feedback in the current study. We also chose to use the participant’s natural remaining sensory circuitry for object touch decoding and address the need of patients with SCI to use their own hand during upper limb activity^9^.

BCIs are emerging as a new means to treat patients suffering from an array of functional deficits^1–8^. Accurately and consistently decoding a single device control signal is a significant challenge for BCIs. Here we extend capabilities of BCI technology to simultaneously decipher multiplexed^12^ afferent and efferent neural activity and dynamically control motor and sensory augmentation devices. Our results support the hypothesis that sub-perceptual residual neural information can be reliably decoded from the human brain, and transformed to conscious perception to augment function.

BCI electrode arrays for treating upper limb dysfunction are almost exclusively implanted in neural tissue to decode motor intention signals alone^1–8^. The sensorimotor demultiplexing capability should impact how BCI electrode array implant locations are determined, for interfaces seeking to decode multiplexed information classes relevant for BCI control. It will be critical to perform multimodal pre-surgical brain mapping to localize these relevant neural representations and further inform electrode array implant location. For example, seemingly small areas of the nervous system may simultaneously encode multiplexed^12^ classes of sensory, motor, and contextual information relevant for next-generation neural interfaces and context adapting sensorimotor therapeutics. We anticipate future efforts that maximize information extraction from neural data, and significantly increase the capability neural interfaces. Furthermore, the data presented here is from a neural interface that has been implanted for over 5 years (at the time of this writing). Reliable next-generation neural interfaces will also need to function for many years to mediate long-term benefits in patients^27,28^.

Human cortex is generally modular and can encode a variety of stimuli or other activity. The sensory signal utilized in this study may arrive in M1 directly, or from a separate source via propagating activity^19^. Furthermore, evidence is accumulating that M1, and other cortical modules, encode a multiplicity of features related to experience beyond their primary processing designation^20–25^. For future BCI applications, an array of powerful control signals can potentially be demultiplexed from a single recording site, or multiple distributed interfaces. Advanced decoding strategies^6^ may be needed to decipher the multitude of representations encoded in neural activity and further enable demultiplexing BCIs. Regardless, the results we present here are a step towards the design of next-generation neural interfaces capable of demultiplexing multimodal neural information for distributed device control and functional improvement.

## Acknowledgements

We would like to thank the study participant for his contributions and exceptional dedication to the ongoing clinical study and technology development. We would also like to thank our development and management teams at Battelle Memorial Institute, Nathan Platfoot and Nick Annetta for helping construct the sensory feedback interface electronics, Colin Dunlap for his assistance in the clinical assessments, Russ Kittel for his contributions to the manuscript graphics, and several Battelle team members who contributed to the preparation and review of the manuscript. A special thanks to Dr. Keith Tansey for his manuscript reviews, and critical insights regarding the clinical importance of these findings. Financial support for this study came from Battelle Memorial Institute and The Ohio State University Center for Neuromodulation.

## Author Contributions

P.D.G., S.C.C., M.A.S., M.A.B., and D.A.F. conceived and designed the experiments. P.D.G., S.C.C., C.E.S., M.A.B., and A.F.J. performed the experiments and analysis. P.D.G., D.J.W., and G.S. provided project supervision. All authors contributed to writing and editing the manuscript.

## Materials & Correspondence

Correspondence and materials requests should be addressed to PDG.

