## Supplemental for "Augmenting Quadriplegic Hand Function Using a Sensorimotor Demultiplexing Neural Interface"

Table of Contents:

A) Methods ………………………………………………………………………….. p. 2 - 15
B) Supplemental Figures
   a. Figure S1………………………………………………………………... p. 16
   b. Figure S2………………………………………………………………... p. 17
   c. Figure S3………………………………………………………………... p. 18
   d. Figure S4………………………………………………………………... p. 19
   e. Figure S5………………………………………………………………... p. 20
   f. Figure S6………………………………………………………………... p. 21
C) Supplemental Video Legends ……………...……………………………………... p. 22
D) Supplemental Appendix References ……………..……………………………. p. 23 - 24

**Methods:**

**Common Methods:**

**i) Study Participant.** The study participant was a 27-year-old male with stable, non-spastic C5 quadriplegia resulting from cervical SCI. As outlined in our previous study[4], the participant had full bilateral elbow flexion (grade 5/5), active wrist extension with radial deviation through an incomplete range of motion against gravity (grade 2/5), but no motor function below the level of C6. His sensory level was C5 on the right (because of altered but present light touch on his thumb) and C6 on the left. His injury was clinically complete, with an overall neurological level of C5 American Spinal Injury Association Impairment Scale A (AIS-A), with zone of partial preservation for motor function to C6 bilaterally according to the International Standards for Neurological Classification of Spinal Cord Injury[42].

The participant underwent implantation of a 96 channel Utah microelectrode recording array (Blackrock Microsystems, Inc.; Salt Lake, Utah) in his left primary motor cortex (Fig. 1D). The hand area of motor cortex was identified preoperatively by fusing functional magnetic resonance imaging (fMRI) activation maps obtained while the patient attempted movements co-registered to the preoperative planning MRI. Full details of the fMRI and surgical procedures can be found in reference 4.

**ii) Neural Recording System.** Neural data was acquired using a Utah microelectrode recording array (Blackrock Microsystems, Inc.; Salt Lake City, Utah) and the Neuroport neural data acquisition system. Recorded data from all 96 recording array channels was sampled at 30 kHz and band pass filtered online from 0·3 – 7·5 kHz using a third order Butterworth analog hardware filter. The neural data was then digitized and sent to a PC for saving or further on-line processing using a custom interface in MATLAB 2014a (The MathWorks; Natick, MA).

**iii) Neural Signal Conditioning and Decoding Using Support Vector Machines.** We used stimulation artifact removal, mean wavelet power (MWP) estimation, and non-linear support vector machine (SVM) decoding, similar to our previous studies[4,6,7,26-28]. Functional electrical stimulation (FES) induced stimulation artifacts were detected by threshold crossings of 500 μV occurring simultaneously on at least 4 of 12 randomly selected channels. A 3·5 ms window of data around each detected artifact was then removed and adjacent data segments were rejoined. This approach leaves the vast majority of the neural data intact. Our series of control experiments confirm the removal of the stimulation artifact across several contexts (specifically Supplemental Fig. S3A & S3B; Fig. 2A, FES alone). Data collected here combined with our previous studies demonstrates the robust ability to remove artifacts from the data with this approach prior to signal analysis.

Neural activity was next measured using MWP, similar to our previous studies[4-7,26-28]. Wavelet decomposition was applied to the raw voltage data, using the 'db4' mother wavelet and 11 wavelet scales. Wavelet scales 3–6 were used, corresponding to the multiunit frequency band spanning approximately 234 to 3,750Hz. The mean of the wavelet coefficients for each scale of each channel was calculated every 100 ms and a 1 s wide boxcar filter was applied to smooth the data. Baseline drift in the data was estimated by using a 15 s boxcar filter and was subtracted from the smoothed mean wavelet coefficients for the corresponding 100 ms window. The mean coefficients were then standardized per channel, per scale, by subtracting the mean and dividing by the standard deviation of those scales and channels during the training blocks. The four scales were then combined by averaging the standardized coefficients for each channel, resulting in 96

MWP values, one for each electrode in the recording array, for every 100 ms of data. The resulting MWP values were used as input to the given non-linear SVM decoder[29,30]. The SVM model training and testing methods are detailed below for both the passive sensory stimulation or active object touch experiments.

**iv) FES system.** The FES system used to stimulate the arm musculature and produce movement was identical to our previous studies[4-7,26-28]. The FES system consists of a multi-channel stimulator and a flexible cuff containing 130 electrodes that is wrapped around the participant's forearm. During use, hydrogel disks (Axelgaard; Fallbrook, CA) were placed between the electrodes and skin to act as a conduction enhancer. The electrodes are 12 mm in diameter and were spaced at 22 mm intervals along the longitudinal axis of the forearm and 15 mm intervals in the transverse direction. Current-controlled, monophasic rectangular pulses (50 Hz pulse rate and 500 μs pulse width) were used to provide electrical stimulation and produce movement. Pulse amplitudes ranged from 0 to 20 mA and were updated every 100 ms. Stimulator calibrations were performed for a given movement using an anatomy-based trial-and-error method to determine appropriate electrode spatial patterns.

<u>All methods below are separated into either passive sensory stimulation or active object touch experiments. All experiments were performed across a total of approximately 1 year.</u>

### <u>Passive Sensory Stimulation Experiments:</u>

#### 1) Passive Sensory Stimulation

We first assessed evoked neural activity in left primary motor cortex M1 using bi-polar electro-tactile stimulation at skin locations on the participant’s arm and hand. Electro-tactile stimulation was chosen for several reasons: 1) its use in our previous FES studies[4-7,26-28]; 2) its safety and precise electronic control of stimulus timing and intensity; and 3) its ability to evoke activity in M1 following from our pilot recordings. We targeted 4 skin locations innervated by the spinal cord above, at, and below the participant's C5 SCI (Fig. 1C). The skin stimulation locations were located at the following dermatomes[32,41]:

1. C5 dermatome (forearm; electrode location: skin above the extensor carpi radialis longus)
2. C6 dermatome (thumb; electrode location: skin above the distal phalanx of digit 1)
3. C6 / C7 dermatome (index; electrode location: skin above the distal phalanx of digit 2)
4. C7 / C8 dermatome (middle; electrode location: skin above the distal phalanx of digit 3)

Forearm, thumb, index, and middle are used to describe these 4 skin stimulation sites throughout the manuscript. A subset of control recordings were also performed on the opposite arm ipsilateral to the M1 implant for the homotopic thumb and forearm locations at the maximum stimulation intensity (Supplemental Fig. S3). Cutaneous landmarks and/or ink markings were used throughout as needed to confirm skin stimulation locations. The participant wore an eye mask and ear plugs during all passive sensory stimulation experiments to significantly reduce any external visual and auditory events during recordings. Recordings were video recorded and performed under the supervision of a licensed physiatrist.

The stimulation interface for a given skin location consisted of a pair of hydrogel disk electrodes adhered to a modified version of the FES interface used in our previous studies[4-7,26-28]. Each hydrogel disk electrode (Axelgaard; Fallbrook, CA) is 12 mm in diameter, 1.27 mm thick, spaced by ~2-3 mm, and attached to a metal electrode consisting of copper with an electroless nickel immersion gold coating embedded in the polyimide flex circuit. We used two current levels of stimulation: minimum intensity = 2·4 mA, and maximum intensity = 9·6 mA (current controlled stimulation, monophasic rectangular pulses, 50 Hz, 500 μs pulse width, 100 ms train duration). Stimulation intensity was selected based on our pilot studies to apply stimulation sufficient to evoke activity in M1 (minimum intensity) and up to an intensity below a noxious level (maximum intensity). At a given skin stimulation location, fifty replicates of stimulation were performed within a given recording with an inter-stimulus interval of 2 s, to ensure the relaxation of neural activity similar to our previous studies[24,25]. On a given recording day, 2 skin locations were selected randomly for stimulation and simultaneous neural recordings. The order of stimulation amplitude for a given skin stimulation location was also selected randomly (e.g., a recording at maximum intensity followed by a recording at minimum intensity, or vice versa). Following a given recording, the participant was asked if he felt the stimuli and whether it was higher or lower in intensity than the previous recording, if applicable (related to Supplemental Fig. S4). We performed a total of 5 recordings at a given skin stimulation site and stimulation intensity. Recordings were performed across a total of ~5 months to assess the chronic viability of the evoked neural signal.

**2) Evoked Activity Analyses**

*Peri-Stimulus Time Histograms (PSTHs)*

All neural recordings were analyzed offline using MATLAB 2016b (The MathWorks). Following stimulation artifact removal (see *Common Methods* above), the signal was band-passed filtered (3$^{rd}$ order Butterworth filter; 300 – 3000 Hz). Multi-unit activity was classified off-line using superparamagnetic clustering (Wave clus)[31]. Unit clusters were manually inspected

(Supplemental Fig. S1A & S1B), similar to our previous studies[24,25]. For each channel, the multi-unit neural activity was first binned (20 ms bin width). We calculated evoked responses on a channel by channel basis (total of 96 recording array channels). A peri-stimulus time histogram (PSTH) was constructed for a given channel using the binned neural data 1 second before and after the start of stimulation, averaged across all stimulation trials (example single channel PSTH: Supplemental Fig. S1C). The magnitude of the evoked response was then quantified similar to our previous studies using PSTH-based analyses[24,25]. For a given channel, a response was considered significant if 1) it exceeded an activation threshold set as the average background activity of the channel across all stimulation trials (evaluated from -1 to -0·02 s before the stimulus) plus three standard deviations (Supplemental Fig. S1C, horizontal gray dashed line), and 2) at least three bins were over the activation threshold. The response magnitude for a given channel was then quantified as the background-subtracted number of spikes within the post-stimulus window of the PSTH (0 – 1 s after the stimulus). The response magnitude for a given channel was zero if it did not meet the significance criteria listed above. The global response magnitude was then estimated using the average response magnitude across all array channels for the given condition. We report the global response magnitude (average spikes per channel; Fig. 1F, Supplemental Fig. S2B). Global PSTHs are background subtracted and smoothed with a 1$^{st}$ degree polynomial model for plotting purposes (Fig. 1E, Supplemental Fig. S2A).

*Decoding Passive Sensory Stimulation*

A nonlinear support vector machine (SVM) classifier was used to decode stimulus location for the passive sensory stimulation recordings at both the minimum and maximum stimulation intensity (referred to as a ‘passive sensory decoder’ in the results) (SVM hyperparameters: $\gamma$ = 0·001, C = 1)[29,30]. A separate SVM model for each stimulation intensity was built with 5 classes:

Rest, Forearm, Thumb, Index, and Middle (related to Fig. 1H & Supplemental Fig. S2C). The input features for each model were calculated as follows: (1) We recorded neural activity and calculated MWP during ~250 total stimuli for each stimulus intensity and skin location across ~5 months; (2) MWP was standardized across blocks within each day to account for day-to-day variability; (3) For each skin stimulation trial, defined by 0·2 s before and 0·8 s after a given sensory stimulus (this epoch was chosen due to the robust neural modulation that occurs during this time period around the stimulus, see Fig. 1B, Supplemental Fig. S2A), MWP was vectorized to a 960-feature vector (96 channels * 10 bins; where each bin spanned 100 ms); The same MWP vectorization process was applied to 250 randomly selected 1 s samples of Rest data collected during spontaneous activity in the first 15 s of a given recording; and (4) For each class, the vectorized MWP was shuffled to remove any effect of stimulus order or recording time during the ~5-month period and equally assigned to either training or testing data (~125 trials per class for training; ~125 trials per class for testing). We report SVM model performance as a confusion matrix (diagonal values = sensitivity; off-diagonal values = false positive rate). Sensitivity is calculated as the percentage of correctly predicted class labels for the targeted class (i.e., true positive rate). False positive rate is calculated as the percentage of incorrectly predicted class labels for a given off-target class. Rows represent the actual recorded class, while the columns represent the model's predictions (related to Fig. 1H & Supplemental Fig. S2C).

### Active Object Touch Experiments:

#### 1) Clinical Assessment of Sensory Function

Monofilament testing was performed by a licensed physiatrist to evaluate the participant's hand sensory function (GRASSP assessment, Semmes-Weinstein monofilaments; Toronto, ON)[32]. All sensory function testing was performed in the absence of visual feedback, as per the

International Standards for Neurological Classification of Spinal Cord Injury published by the American Spinal Injury Association[26]. The palmar and dorsal aspects of digits 1 (thumb), 3 (middle), and 5 (pinky) were exposed to multiple trials of either 0·4, 2, 4, or 300 g of force while the participant was blind-folded (related to Fig. 1A). Trial location and force level were randomized. The participant was asked to report the application of the applied tactile stimulus. The following scores were generated to quantify the participant's tactile acuity[32]: 4 = 0·4 g detection at 66 %; 3 = 2 g at 33 %; 2 = 4 g at 33 %; 1 = 300 g at 33 %; 0 = 300 g at 0 %.

The participant uses his hand to manipulate objects during BCI operation. We assessed the participant's ability to detect object touch during FES-mediated and grip (standardized objects tested from the Action Research Arm Test[33]: small cylinder (1 cm diameter) and large cylinder (2 cm diameter)). The participant was again blind-folded, and the object was placed between digits 1 and 2 without touching the skin on randomized trials where a grip was triggered (small cylinder: lateral pinch grip; large cylinder: *can* grip). A grip was activated for a duration of 3 seconds for a given trial. The participant then reported whether there was an object in his hand. Each grip trial was bounded by rest periods with random durations between 5 to 6 seconds. We report Object Touch Detection as the percentage of trials the participant correctly identified there was an object present during grip (related to Fig. 1B).

**2) Decoding Active Object Touch**

We trained SVM decoders to recognize active object touch events in real-time (referred to as a 'touch decoder' in the results) (specifically related to Fig. 2 & 3). These decoders were trained using neural data during active object touch, in contrast to the passive sensory decoders described above (see *Decoding Passive Sensory Stimulation*). We used the *can* object, a part of the standard clinical grasp and release test battery (5·4 x 9·1 cm)[6,7,13]. For model training, we recorded 9 total cues of labeled touch data, with each cue consisting of a 6 second period. Cues were conveyed by a virtual hand on a computer monitor. Each cued period of touch data was bounded by rest cues with random durations between 5 to 6 seconds. For each touch cue period, the participant first moved his hand down onto and around the *can* object for 3 seconds, followed by a scripted object grip period for an additional 3 seconds where FES triggered a more forceful grip. Therefore, touch decoder model training consisted of neural data during: 1) movement onto the object, 2) touch of the object, and finally 3) additional FES mediated touch. This touch decoder model was then tested on 4 cue types and rest periods to assess model performance during 'touch' and 'no touch' events. The participant completed the following cued events: (1) 3 seconds of natural touch of the object followed by 3 seconds FES mediated touch ('Touch'), (2) 6 seconds of natural object touch ('Touch'), (3) 6 seconds of identical movement without the object present ('No Touch'), (4) 6 seconds of FES without the object present ('No Touch'). Rest periods were also assessed, and consisted of 5 second epochs randomly selected across all recordings. For this touch decoder testing, we report model responsiveness during the 4 cue types and rest, defined as the percentage of time the touch decoder output was above the activation threshold during the given period (activation threshold = 0·5; see Supplemental Video 1 for exemplary touch decoder outputs during testing). This touch decoder was then used to trigger the closed-loop sensory feedback interface during the object manipulation experiments described below (see *Functional Improvement Assessments With and Without Closed-loop Sensory Feedback* section). In a subset of experiments, we also assessed touch decoder timing during simultaneous recording of applied force (related to Supplemental Fig. S5; force transducer interface: custom designed piezoresistive sensor pad (FlexiForce; Boston, MA) interfaced with an Arduino Mega 2560 board transferring force data to the PC).

#### 3) Decoding Motor Intention

Similar to our previous studies, we built motor decoders for manipulating the *can* object (a part of the grasp and release test)[6,7,13]. Briefly, the participant was prompted to imagine performing a *can* grip and movement, using a virtual hand displayed on a computer monitor. Each motor cue lasted 3-4 seconds, and was bounded by rest cues with random durations between 5 and 6 seconds. During the initial motor cues, FES triggered the *can* grip. FES was controlled by the SVM motor decoder starting on the 4$^{th}$ motor cue. This motor decoder model was updated during subsequent training cues until a sufficiently accurate model was built (accuracy > ~80%). This motor decoder was then used to control FES during the object manipulation experiments described below (see *Functional Improvement Assessments With and Without Closed-loop Sensory Feedback* section).

#### 4) Sensory Feedback Interface

The sensory feedback interface consisted of 3 low-noise vibrotactile coin motors affixed to a velcro band wrapped around the participant's right bicep (cartoon schematic in Fig. 2B & 3A) (coin motor details: 12 mm diameter, 3·4 mm height, 2·6 G force output; Need for Power; Shenzhen, Guangdong, China). This interface was tethered to an Arduino Mega 2560 board to power and control vibrotactile sensory feedback (all 3 vibrotactile motors were either off, or turned on by the touch decoder at their max level: 2·6 G force). Sensory feedback interfaces targeting the skin over the biceps have been used in several sensory feedback studies and is well studied[34-37]. Our pilot data confirm that the participant's right bicep exhibited normal sensory function, and vibrotactile stimulation was recognized on 100% of stimuli. The interface was designed to ensure participant comfort during movement. The vibrotactile motors achieved maximum amplitude within 1 ms of controller signal initiation. All sensory feedback interface communication was also recorded. This sensory feedback interface was controlled by the touch decoders outlined above and was triggered in real time during the closed-loop sensory feedback tasks described below.

**5) Functional Improvement Assessments With and Without Closed-loop Sensory Feedback**

We assessed upper limb function across a battery of 4 clinical assessments under the supervision of a licensed physiatrist. The sensory feedback interface was placed on the participant's right bicep during all assessments, and function was assessed across trials during either a 'Control' or 'Demultiplexing With Sensory Feedback' condition. The 'Control' condition consisted of functional testing without any vibrotactile sensory feedback. 'Demultiplexing With Sensory Feedback' consisted of on-demand touch decoder controlled sensory feedback for rapidly conveying hand touch events back to the user. The participant was blinded to the given condition before a series of assessment trials. All clinical assessments were performed across 2 clinical testing days. The trial counts and statistical tests are described in the *Data Analysis and Statistics* section.

The first clinical assessment was an extension of the monofilament testing described above (see *Clinical Assessment of Sensory Function* section)[32]. A touch decoder was first constructed identical to that described above (see *Decoding Active Object Touch* section). The palmar aspects of digits 1, 3, & 5 were next exposed to multiple trials of either 0·4, 2, 4, or 300 g of force using the monofilaments, in the absence of visual feedback[26]. Trial location and monofilament force level were randomized. This allowed us to map what hand dermatomes contributed to the activation of the touch decoder (digit 1 (thumb): C6; digit 3 (middle): C7 / C8; digit 5 (pinky): C8)[32,41]. We assessed whether the touch decoder was activated following the application of a given stimulus using simultaneously recorded high-speed video. We report the lowest dermatome level the touch decoder was activated by during the monofilament assessment (e.g., palmar tactile stimulation to digits 3 & 5 repeatedly activated the touch decoder (dermatomes innervated by spinal level C7 / C8)).

The second clinical assessment was an extension of the Object Touch Detection test described above (see *Clinical Assessment of Sensory Function* section). The standardized large cylinder object was used[33]. A touch decoder was first constructed identical to that described above (see *Decoding Active Object Touch* & *Decoding Motor Intention* section). The participant was again blind-folded, and the object was placed between digits 1 and 2 without touching the skin on randomized trials. Grip was triggered by FES during a shuffled series of 'Demultiplexing With Sensory Feedback' or 'Control' condition cues. Grip was activated for a duration of 3 seconds for a given trial. The participant then reported whether there was an object in his hand. Each grip trial was bounded by rest periods with random durations between 5 to 6 seconds. We again report Object Touch Detection as the percentage of trials the participant correctly identified there was an object present during grip (related to Fig. 2C).

The third clinical assessment consisted of the modified grasp and release test (GRT) only using the *can* object[13]. A touch decoder and motor decoder was constructed identical to that described above (see *Decoding Active Object Touch* & *Decoding Motor Intention* section). The participant was then cued to repeatedly grasp, move, and release the object during shuffled series of 'Demultiplexing With Sensory Feedback' or 'Control' condition trials. After each GRT trial, the participant reported his sense of agency (SoA) (i.e., "How in control did you feel of the movement and grip?"). The SoA score ranged from 0-100, similar to previous studies[38,39] (0 = poor sense of control; 100 = perfect sense of control; related to Fig. 3C).

The last clinical assessment was a modified GRT again using only the *can* object (related to Fig. 3D). A touch decoder and motor decoder was constructed identical to that described above (see *Decoding Active Object Touch* & *Decoding Motor Intention* section). The participant was instructed to repeatedly grasp, transfer, and release the *can* object onto an elevated platform as fast as possible during shuffled series of ‘Demultiplexing With Sensory Feedback’ or ‘Control’ condition trials. Each GRT assessment period consisted of two 60 second object transfer periods separated by a 20 second rest period. All GRT trials were recorded with high speed video for offline analysis. We quantified the number of objects successfully transferred and the object transfer times, similar to our previous studies[6,7]. We also assessed the interval between the touch decoder and motor decoder activations to examine the neurophysiological substrates of GRT performance with and without sensory feedback (high-speed video was also used in addition to decoder times to confirm touch and motor event start times). The touch decoder or motor decoder start times were calculated across GRT trials using the time each decoder crossed the device activation threshold (device activation threshold = 0·5). We report the interval (s) between the touch and motor decoder activations across testing conditions.

**Data Analysis and Statistics.** Normality tests were performed for each analysis to determine if parametric or nonparametric statistics should be used. All statistical tests were two-tailed unless otherwise noted, and performed in MATLAB 2016b. An alpha level of 0·05 was accepted for significance unless Bonferroni corrections are noted.

Effects of sensory stimuli on evoked M1 neural activity across skin locations were evaluated using separate one-way ANOVAs for the maximum (Fig. 1F) and minimum (Supplemental Fig. S2B) stimulation intensities. The factor was skin location with 4 levels:

forearm, thumb, index, and middle. Tukey's post-hoc test was used to determine differences in global response magnitude across skin locations. Independent samples t-tests were used to determine the effects of sensory stimuli on evoked M1 neural activity for data recorded during stimulation of the contralateral and ipsilateral forearm and thumb (related to Supplemental Fig. S3).

A one-tailed independent samples t-test was used to determine if decoder performance values were above chance for the passive sensory stimulation data (confusion matrices, Fig. 1H & Supplemental Fig. S2C). Each confusion matrix value was compared to a chance prediction level for statistical evaluation. Chance levels were generated by randomly permuting the data labels 10 times[40]. Values from unrandomized label permutation were then compared with values from randomized label permutation. A Bonferroni corrected alpha value of 0·002 was used for significance (0·05 / 25 comparisons). Differences in perceptual and decoder sensitivities were assessed using separate one-way ANOVAs for each of the 4 skin locations (related to Supplemental Fig. S4).

For the active object touch experiments, touch decoder responsiveness values were assessed using a one-way ANOVA. The factor was cue type with 4 levels: Object Touch & FES, Object Touch, FES alone, and Movement alone. Tukey's post-hoc test was used to determine differences in touch decoder responsiveness across cue type. Functional improvement assessments were performed across 2 separate clinical testing days for the following total trial counts: object touch detection: 16 trials (for either the 'Demultiplexing With Sensory Feedback' and 'Control' conditions), SoA: 24 trials (for either 'Demultiplexing With Sensory Feedback' and 'Control' conditions), modified GRT performance & decoder interval: 78 trials ('Demultiplexing With Sensory Feedback') and 72 trials ('Control'). Effects of closed-loop sensory feedback were assessed using independent samples t-tests for the object touch detection, SoA, GRT performance, and decoder interval data, comparing the ‘Demultiplexing With Sensory Feedback’ to ‘Control’ conditions.

**Supplemental Figures:**

**Supplemental Figure S1:**

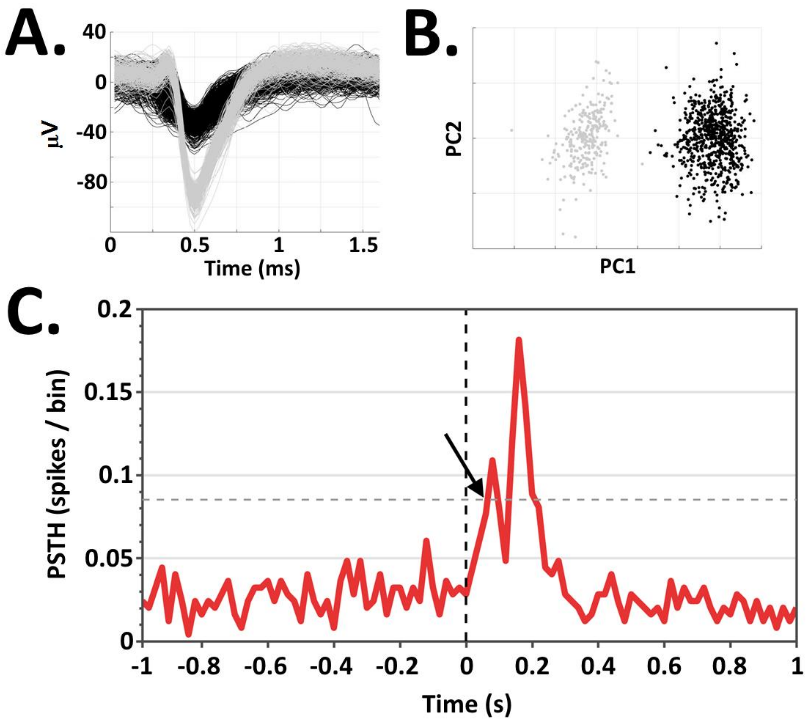


**Supplemental Figure S1. Neural Recording Methods.** Neural activity was recorded from a Utah microelectrode recording array implanted in left M1 (Fig. 1D) during skin stimulation protocols (**A** = exemplar unit waveforms; **B** = corresponding principal component analysis of the unit waveforms). **C.** For each array channel, we constructed a peri-stimulus time histogram (PSTH) of the multiunit activity to assess neural modulation evoked by skin stimulation. Exemplar single channel PSTH is plotted 1 s before and 1 s after the start of skin stimulation (stimulation occurs at time 0, vertical black dashed line). To quantify the neural response, an activation threshold was first calculated based on the background neural activity (activation threshold = horizontal gray dashed line). At least 3 total bins crossed the activation threshold to be considered an evoked response (black arrow: first significant crossing), similar to our previous studies[24,25]. See *Peri-Stimulus Time Histograms* section of the Methods for additional data processing details.

**Supplemental Figure S2:**

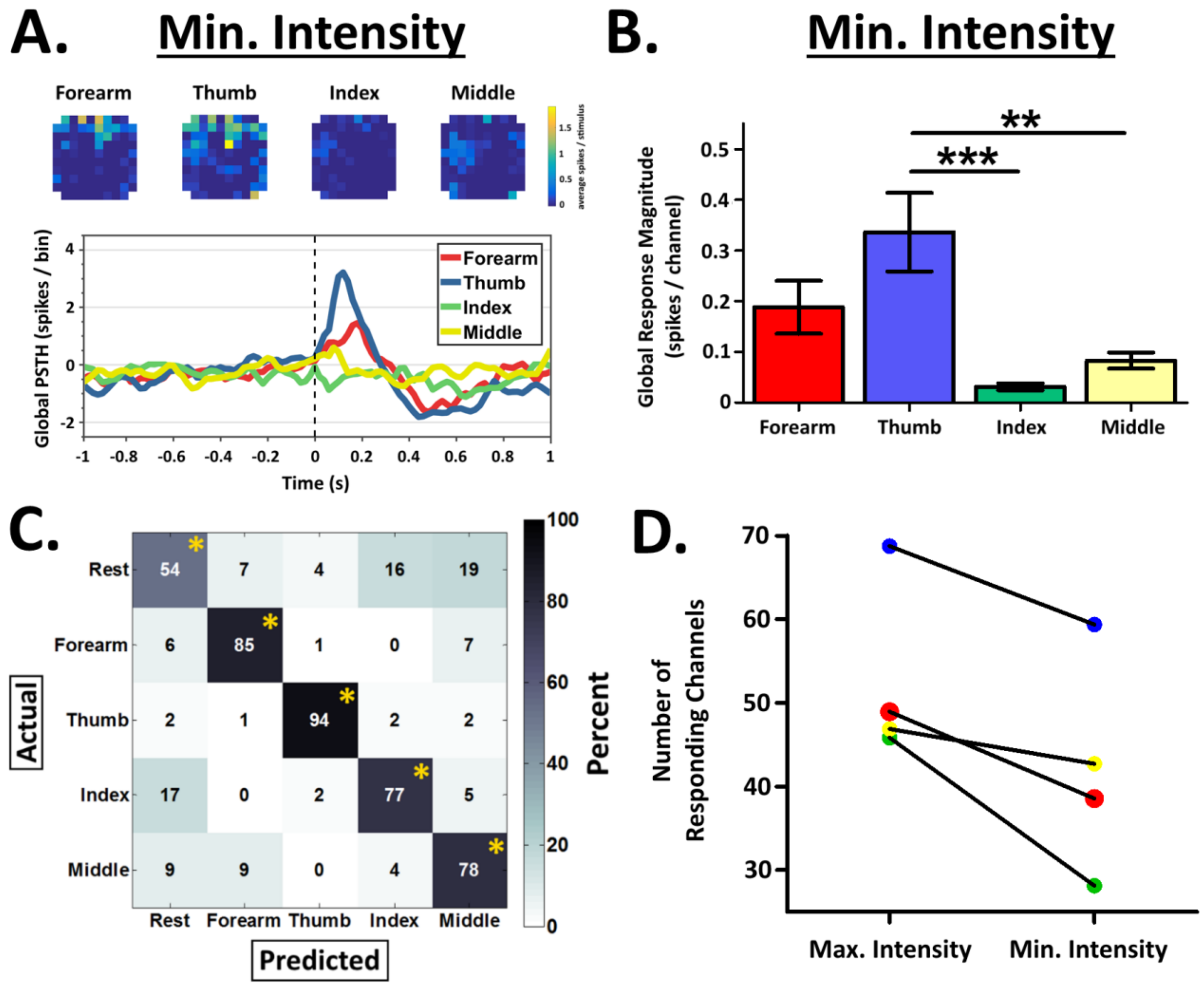


**Supplemental Figure S2. Evoked Multiunit Responses Across Stimulation Intensities. A.** Color coded representations of multiunit response magnitudes across the microelectrode recording array (color scaling: blues = no or small M1 responses; yellow = large M1 responses; units: average spikes / stimulus), and accompanying peri-stimulus time histogram (PSTH) during stimulation to the forearm (red), thumb (blue), index finger (green), or middle finger (yellow) at the minimum stimulation intensity. **B.** At the minimum stimulation intensity, stimulation of the thumb evoked a significantly larger global response magnitude compared to index or middle (F[3, 380]=7·9, p<0·001; *** = p<0·001, ** = p<0·01). **C.** Number of channels responding to the maximum and minimum intensities across skin locations.

**Supplemental Figure S3:**

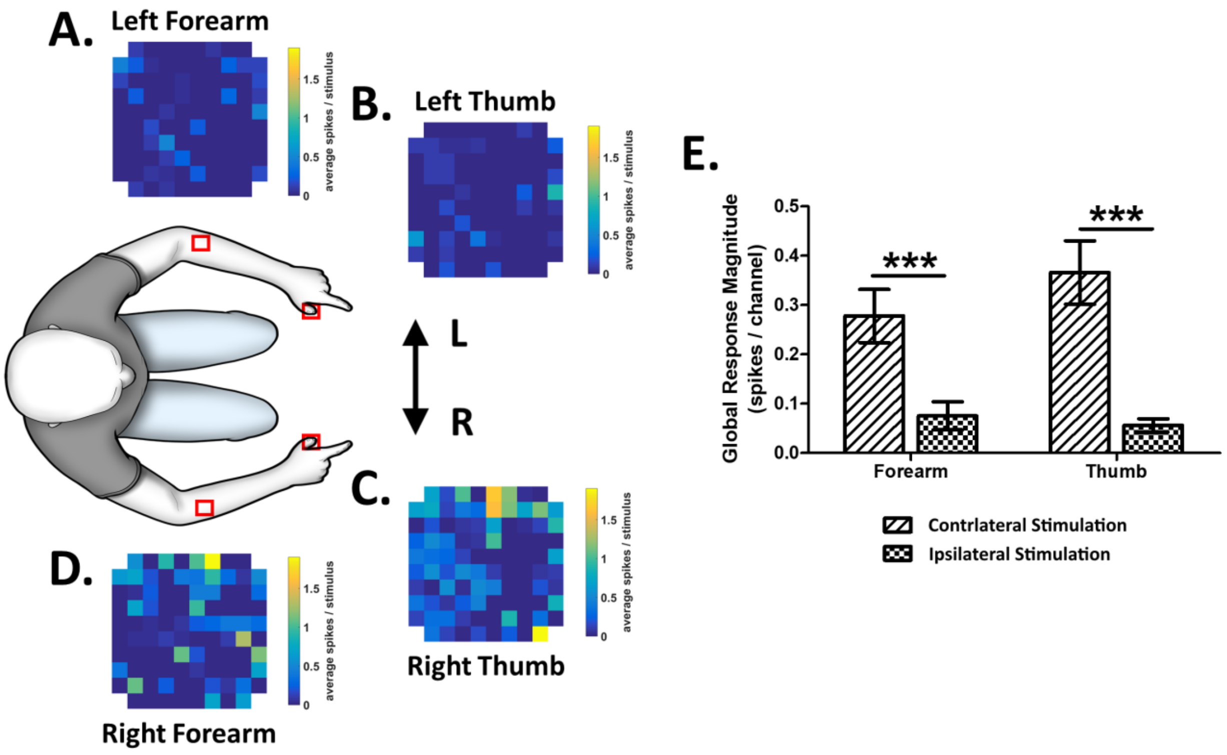


**Supplemental Figure S3. Control Sensory Stimuli Ipsilateral to the Microelectrode Recording Array in Left M1 Evoke Little to No Neural Activity.** We performed control recordings using electrotactile stimuli to the homotopic forearm (**A**) and thumb (**B**) locations on the arm ipsilateral to the M1 recording array at the maximum stimulation intensity (color coded representations of multi-unit response magnitudes are shown across the microelectrode recording array; color scaling: blues = no or small M1 responses; yellow = large M1 responses; units: average spikes / stimulus). As expected, stimuli to these skin sites demonstrate little to no evoked neural activity in left M1. Global response magnitudes following contralateral skin stimulation (at the contralateral thumb (**C**) and forearm (**D**), repeated from Fig. 1F & 1G) were on average over 5 times larger compared to responses following homotopic ipsilateral skin stimulation (**E**, forearm: $t(95) = 3{\cdot}9$, thumb: $t(95) = 4{\cdot}9$; *** = $p<0{\cdot}001$). These results support the hypothesis that evoked activity in M1 following sensory stimuli is maximal from semi-intact skin locations contralateral to the microelectrode recording array. Stimulated skin locations outlined in red boxes.

**Supplemental Figure S4:**

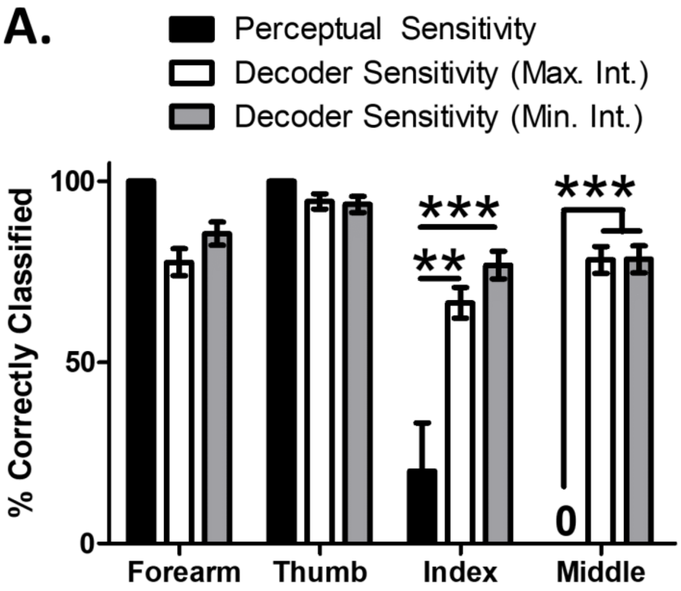


**Supplemental Figure S4. Passive Sensory Decoders Accurately Decipher Sub-perceptual Neural Activity.** We also assessed the participant's perceptual sensitivity (related to the data reported in Fig. 1H & Supplemental Fig. S2C). We defined perceptual sensitivity as the ability of the participant to decipher whether stimulation was simply felt during a given train of stimuli (at the minimum or maximum intensity). As expected, the participant was able to feel stimuli to skin innervated from above or at the level of the SCI at a high rate (forearm and thumb), and was largely unable to feel stimuli to skin innervated from below the level of the SCI (index and middle). Interestingly, passive sensory decoder sensitivities were significantly higher compared to the participant's perceptual sensitivity for stimuli he largely cannot feel (index, $F[2,257] = 8{\cdot}1$, $p < 0{\cdot}001$; middle, $F[2,256] = 17{\cdot}8$, $p < 0{\cdot}001$; ** = $p < 0{\cdot}01$, *** = $p < 0{\cdot}001$; decoder sensitivity values repeated from Fig. 1H & Supplemental Fig. S2C). These results demonstrate the ability to decode residual sensory neural activity that is largely below conscious perception. Data presented are mean ± S.E.M.

**Supplemental Figure S5:**

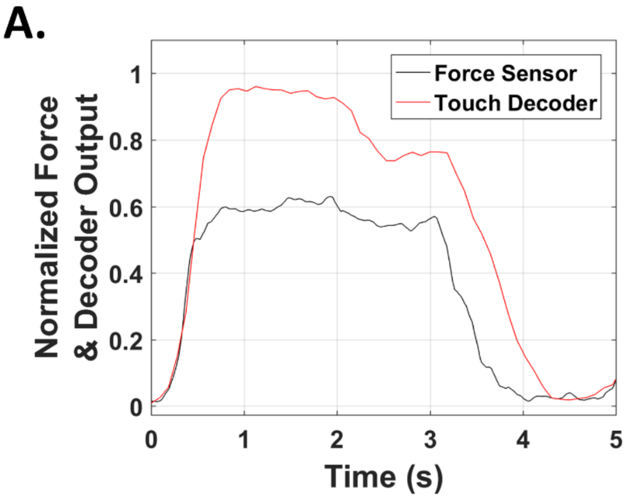


**Supplemental Figure S5. Touch Decoders During Object Grip Are Time Locked to Force Application.** In a subset of experiments, we assessed touch decoder outputs and subsequent force generation during object grip. We applied preprogrammed FES to generate a lateral pinch grip, subsequently creating force transduced by a piezoresistive sensor interface. Neural activity, touch decoders, and force sensor readings were all recorded simultaneously. **A.** The average latency between touch decoder activation and force application was 22 ms. Touch decoder activation was therefore time-locked to force generation, with synchronized on and off times.

**Supplemental Figure S6:**

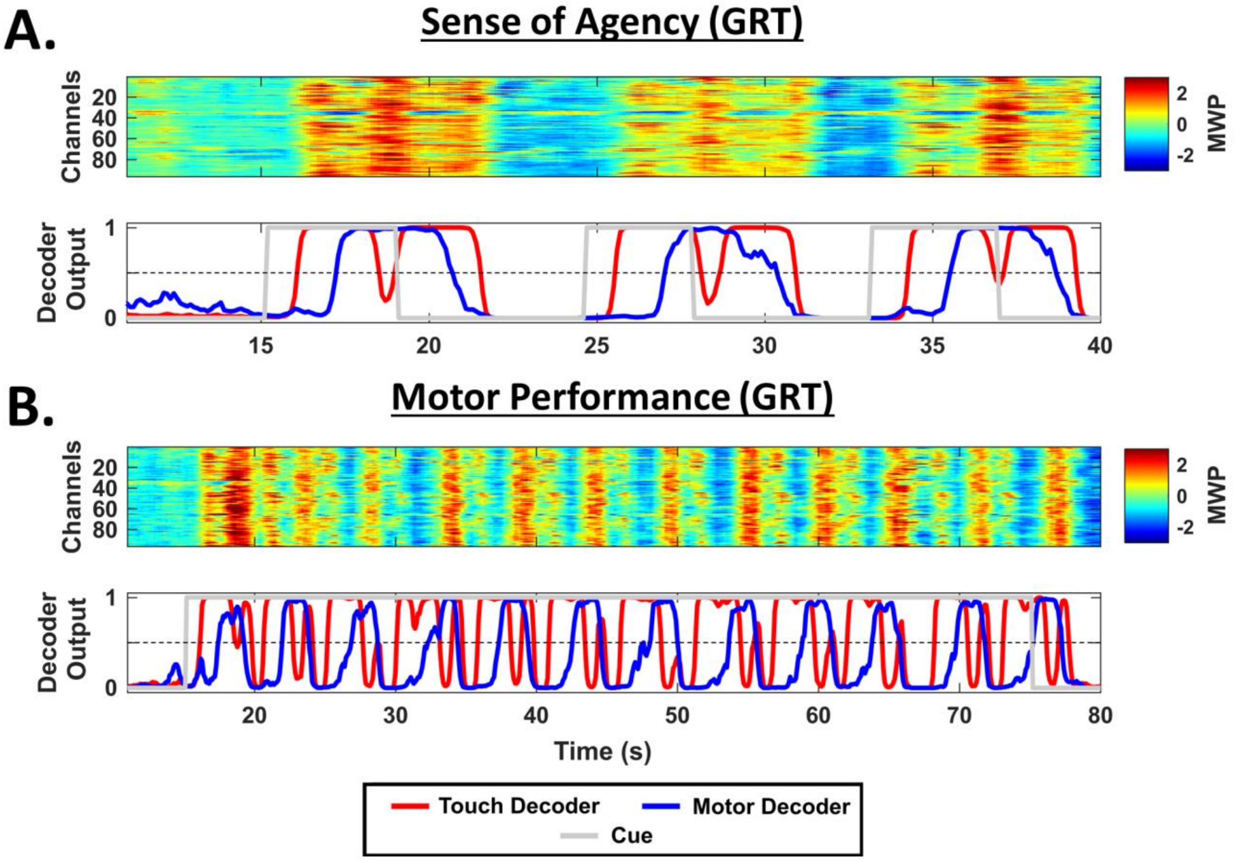


**Supplemental Figure S6. Mean Wavelet Power Input and Decoder Output Plots for the 'Sensorimotor Demultiplexing' BCI Tasks.** Exemplary MWP input and SVM decoder outputs during the sense of agency (**A**; main data: Fig. 3C), and GRT motor performance assessments (**B**; main data: Fig. 3D). Color coded MWP data recorded across the 96-channel microelectrode recording array is shown at the top of each panel, and serve as an input to the SVM. Simultaneous SVM decoder outputs are shown below each MWP plot. Touch decoder outputs (red lines) control the sensory feedback device. Motor decoder outputs (blue lines) control FES of the arm. The device activation threshold is shown as a horizontal dashed line on all decoder output plots (at 0·5). Assessments are cued (overlaid cue periods in gray). Please see the *Active Object Manipulation Experiments* section of the Methods for further details.

### Supplemental Video Legends

**Supplemental Video 1 (Related to Fig. 2A).** The touch decoder was tested during 'touch' and 'no touch' cue types (see *Decoding Active Object Touch* of the Methods for more details). Video contains the cue type (left most column), video of the participant's hand and arm, and the synchronized touch decoder outputs during the given cue type (red trace = touch decoder output; overlaid gray epochs = cue periods; horizontal dashed line = activation threshold). **A.** The touch decoder was robustly activated during 'touch' cues. **B. & C.** Control 'no touch' cues did not activate the touch decoder ('Rest' occurs when the cued period is off). Data shown are exemplary cues from data presented in Figure 2A.

**Supplemental Video 2 (Related to Fig. 3B).** Sensorimotor demultiplexing was performed using separate support vector machine models trained to recognize touch events (touch decoder; red trace in bottom right panel) or motor intention events (motor decoder; red trace in bottom right panel). 'Cortex' panel shows the pseudocolored surface of the patient's brain with the recording array location in the black box (adapted from Figure 1D; S1 = primary somatosensory cortex, M1 = primary motor cortex, PMC = pre-motor cortex). Live color-coded mean wavelet power (MWP) data is shown at the top right, synchronized to the video of the participant's hand and object shown at the left of the video. The participant was repeatedly cued to position his hand around the *can* object, and then generate motor intention to activate FES and transfer the object. Touch decoders were time-locked to object touch. As expected, motor decoders followed the touch decoders during each cued transfer (horizontal dashed line = activation threshold).
